# Emergence of social inequality in a spatial-ecological public goods game

**DOI:** 10.1101/412700

**Authors:** Jaideep Joshi, Åke Brännström, Ulf Dieckmann

## Abstract

Spatial ecological public goods, such as forests, grasslands, and fish stocks risk being overexploited by selfish consumers, a phenomenon called “the tragedy of commons”. The spatial and ecological dimensions introduce new features absent in non spatio-ecological contexts, such as consumer mobility, incomplete information availability, and rapid evolution by social learning. It is unclear how these different processes interact to influence the harvesting and dispersal strategies of consumers. To answer these questions, we develop and analyze an individual-based, spatially-structured evolutionary model with explicit resource dynamics. We find that, 1) When harvesting efficiency is low, consumers evolve a sedentary harvesting strategy, with which resources are harvested sustainably, but harvesting rates remain far below their maximum sustainable value. 2) As harvesting efficiency increases, consumers adopt a mobile ‘consume-and-disperse’ strategy, which is sustainable, equitable, and allows for maximum sustainable yield. 3) Further increase in harvesting efficiency leads to large-scale overexploitation. 4) If costs of dispersal are significant, increased harvesting efficiency also leads to social inequality between frugal sedentary consumers and overexploitative mobile consumers. Whereas overexploitation can occur without social inequality, social inequality always leads to overexploitation. Thus, we identify four conditions, which are characteristic (and as such positive) features of modern societies resulting from technological progress, but also risk promoting social inequality and unsustainable resource use: high harvesting efficiency, moderately low costs of dispersal, high consumer density, and consumers’ tendency to rapidly adopt new strategies. We also show that access to global information, which is also a feature of modern societies, may help mitigate these risks.

## 1 Introduction

A public good that is freely accessible to everyone, but limited in quantity, can be optimally used only if everyone cooperates in using no more than their fair share. However, those who consume more than their fair share obtain a greater benefit. This creates an incentive to overexploit the public good, which threatens to reduce resource availability, leaving everyone worse off. This social dilemma is called the ‘tragedy of the commons’ [1, 2]. The ubiquity of social dilemmas has led to widespread interest in exploring mechanisms that can support cooperation, not only in evolutionary biology but also in economics, cognitive sciences, and social sciences [3–7]. Indeed, research into mechanisms capable of preventing social dilemmas can help inform important policy decisions, such as fisheries regulations, and support treaty negotiations on global commons, such as ozone-layer protection and climate change mitigation [8–17].

Owing to the combined efforts of many scientists, several mechanisms that promote cooperation have been identified [see e.g., 18]. Kin-selection [19], in which cooperative acts are directed towards genetic relatives, and direct reciprocity [20–24], in which individuals cooperate with individuals who return cooperation, are two key mechanisms that promote cooperation. In the case of humans, indirect reciprocity, in which individuals cooperate with those who have a reputation for cooperating [25–27], is also an important mechanism supporting cooperation. Additionally, combinations of positive and negative incentives, such as offering rewards to cooperators and punishing defectors can also be a powerful means of promoting cooperation [28–31]. Such studies are mostly stylized, often based on simple mathematical models and occasionally on experiments.

Most real socio-ecological systems are spatially structured. With some exceptions [e.g., 32], spatial structure is thought to promote cooperation as it both exposes defectors to the consequences of their own selfish acts and allows local clusters of cooperators to form, enabling multilevel selection for cooperation [33–35]. However, space also allows mobility. Mobility can hinder cooperation allowing defectors to escape the consequences of their own acts [36]. Indeed, several studies have considered the effect of fixed mobility and found that cooperation can be sustained when mobility is either low [37] or dependent on local conditions [38, 39]. If mobility incurs no cost, then defectors may invariably evolve high mobility and undermine cooperation. When dispersal is costly, an evolutionary interplay between mobility and cooperation may occur, but thus far only a handful of studies have considered the joint evolution of costly mobility and cooperation [40–44]. A recent study by Mullon et al. [45] found that when dispersal and cooperation coevolve, two coexisting strategies can spontaneously emerge: one benevolent and sessile, the other self-serving and dispersing.

In socio-ecological settings, the ecological dynamics of the resource plays a crucial role in the evolutionary dynamics. First, interactions between individuals are often mediated through the resource, i.e., individuals do not directly interact with other individuals, but respond to changes in the resource caused by others. Second, limited resource availability may lead to the evolution of density-dependent strategies, since per-capita resource extraction is expected to decline with consumer density. Third, ecological public goods, such as renewable resources, have their own ecological time scale of resource replenishment. Most studies assume that evolution is a slow process. However, memetic evolution is not biologically constrained, and can occur on ecological or even faster timescales. Evolutionary outcomes may be dramatically different depending on whether evolution is slow or fast [46]. Furthermore, spatial extent may prevent consumers from having full information about other consumers and the environment, because information may not reach far-off consumers. Such local information may lead to local selection, which is typically expected to benefit defectors. Despite these important gaps, only a few studies have so far considered ecological public goods [2, 47, 48], and even fewer studies explicitly model a renewable resource [e.g., 49].

Here, we move beyond the aforementioned studies, in particular the work by Parvinen [44] and Mullon et al. [45], by investigating the role of explicit resource dynamics, fast evolution, and incomplete information about other individuals’ strategies on the evolution of coopeartion. Using a socio-ecological model of public-goods utilization, we investigate for the first time the coevolution of resource harvesting and dispersal strategies in realistic socio-ecological settings.

## 2 Methods summary

We model consumers and resources on a continuous two-dimensional space. We assume that resource growth is logistic with intrinsic growth rate *r* and carrying capacity *K*. Each consumer *i* exploits resources in his/her local neighborhood at a harvesting rate *r*_H,*i*_. When the local resource falls below a threshold, consumers disperse to new locations drawn from a normal distribution centered at their current position with a mean ‘dispersal radius’ of *σ*_D,*i*_. The harvested resource *R*_*i*_ yields benefit *bR*_*i*_ and incurs cost 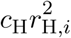, while dispersal incurs cost *c*_D_ per unit length. The payoff to each individual is the time-averaged utility of resource harvest 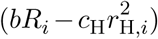 minus the cost of total dispersal distance.

To account for evolution by social learning, we assume that at rate *r* _I_, individuals imitate the strategies (harvesting rate and dispersal radius) of other individuals with greater payoff. The imitation rate defines the evolutionary timescale: consumers are ‘impatient’ if the imitation rate is much larger than the resource growth rate; such consumers tend to imitate even before the consequences of others’ strategies on the resource become apparent. To account for information availability, we assume that each individual is more likely to choose an individual to imitate within an ‘imitation radius’ *σ* _I_. A low imitation radius means that consumers are ‘myopic’ and only imitate their neighbors, whereas a very large or infinite imitation radius means that consumers have ‘global information’, and can imitate anyone in the population. In this way, we model the essential characteristics of human behavior to make our model realistic, while leaving out other confounding factors to keep our model minimal.

Without loss of generality, we reduce the number of parameters by carefully choosing our units. Full details of our model and its parameterization are given in Methods. SI-Fig 1 shows a schematic of the system behaviour along with meanings of key parameter values. Video S5.1 (see https://github.com/jaideep777/jaideep_thesis_videos) shows a representative simulation of consumer behaviour where they exploit local resource and disperse.

**Figure 1:**
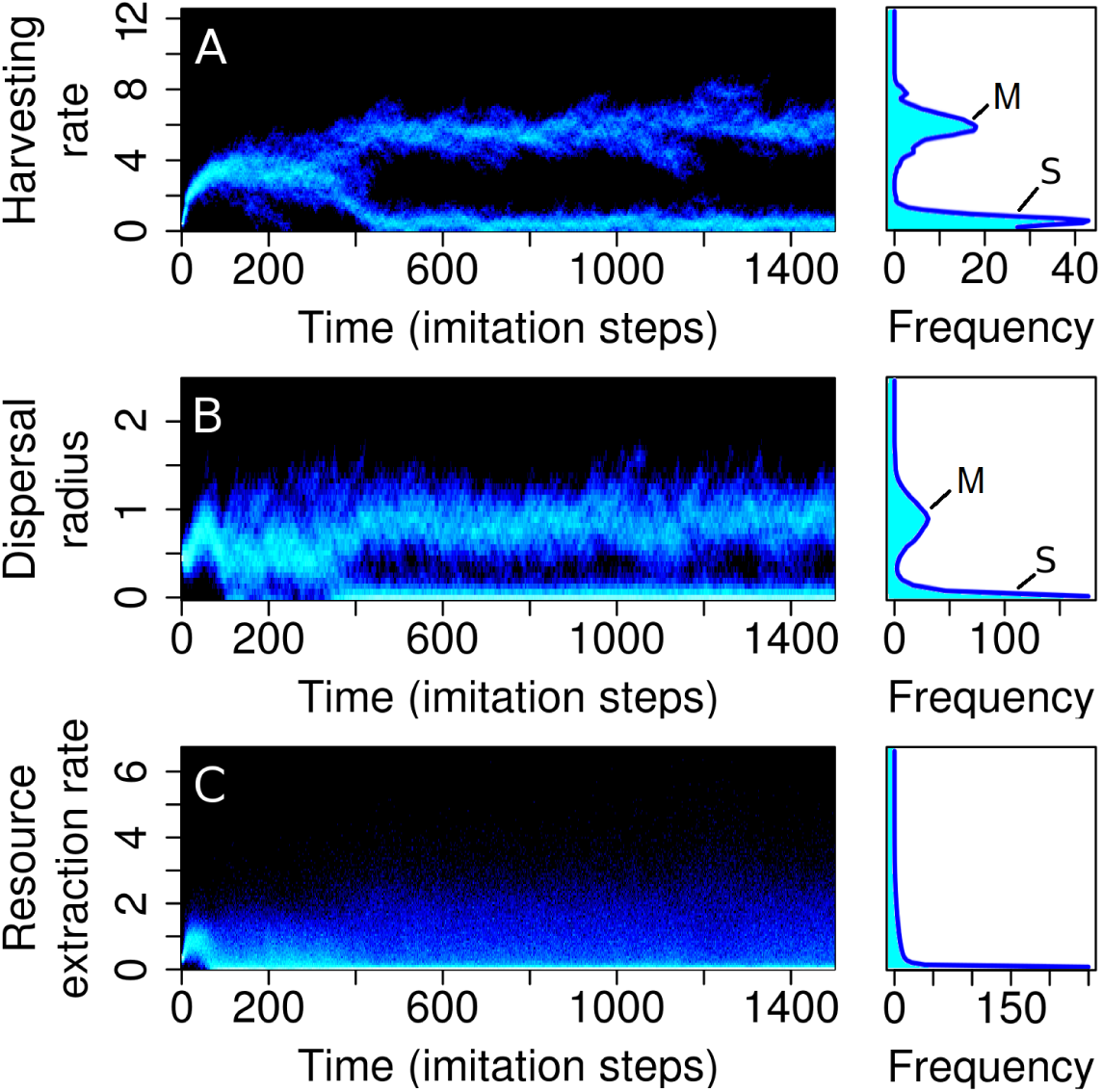
Sedentary and mobile strategies emerge spontaneously. Distributions of harvesting rate (A), dispersal radius (B), and resource extraction rate (C) in the population over time, shown alongside the time-averaged (evolved) strategy distributions. Brighter colors indicate higher frequency, but note that the color scale is non-linear. The population starts with all individuals having identical strategies. As time progresses, social learning leads to spontaneous emergence of two distinct strategies: a sedentary strategy with a low harvesting rate and zero dispersal, and a mobile strategy with a high harvesting rate and high dispersal. Parameters: *b* = 0.059, *c* = 0.68, *r* _I_ = 0.1. Other parameters are as in Table 1.

An important parameter in our model is the consumer density *ρ*. Additionally, our model has two pairs of key parameters. The first pair are the payoff parameters *b* and *c* _H_ that determine the efficiency *b*/*c*_H_ and utility 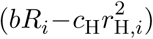 of resource consumption. The second pair are imitation parameters *r* _I_ and *σ*I that determine the spatio-temporal scales of imitation, reflecting impatience and myopia of consumers. See SI-Table 2 for a glossary of terms.

## 3 Results

To explore the social dynamics of harvesting and dispersal in a realistic environment, we perform simulations of our system for a wide range of parameter values, which corespond to diverse socio-ecological settings. Specifically, we investigate the effect of population density, costs and benefits of harvesting and dispersal, the imitation rate, and imitation radius, on the evolved strategies. We present our key findings below.

### 3.1 Sedentary and mobile strategies emerge spontaneously

All individuals start with the same harvesting rate and dispersal-kernel size at time *t* = 0. As the interplay of resource and imitation dynamics progresses, individuals diversify into two distinct strategies (Fig. 1). One is a sedentary strategy, with consumers who adopt a low harvesting rate (*r*_H_ = 0.5) and do not disperse. The second is a mobile strategy, with consumers who adopt a high harvesting rate (*r*_H_ = 6) and a non-zero dispersal radius (*σ* _D_ = 1 in the outcome shown). At the population level, these strategies coexist over time, although individuals may change their strategy frequently by imitation of other individuals.

To investigate the robustness of this result under more realistic assumptions of intelligent consumers, we allow consumers to ‘explore’ random strategies at a low rate. SI-Fig. 2 shows that with low strategy exploration rates, the system evolves to the same evolutionary endpoints as in Fig. 1.

**Figure 2:**
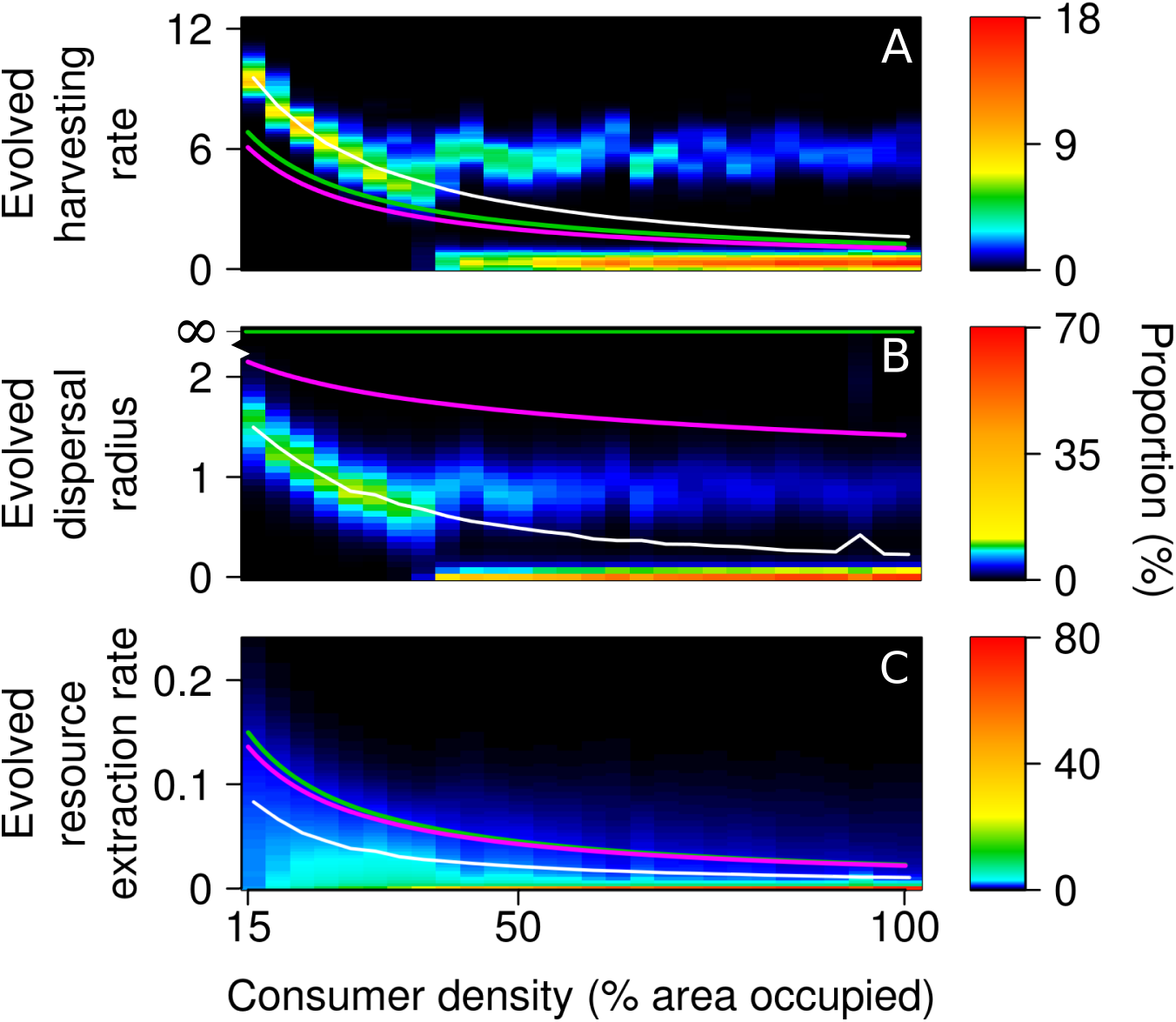
Increasing consumer density leads to social inequality. When consumer density is low, all consumers harvest at high rates and disperse, because available resources suffice for everyone. As consumer density increases, everyone harvesting at high rates becomes unsustainable, leading to diversification of strategies into prudent sedentary consumers and overexploitative mobile consumers (A-B). Average per capita resource extraction rate (% of total carrying capacity of the system) decreases with density (C). However, the average harvesting rate (white line) is higher than the corresponding yield-maximizing (green line) and profit-maximizing (magenta line) rates, leading to overexploitation of resource and suboptimal per capita resource extraction. Parameters: *b* = 0.059, *c* = 0.68, *r* _I_ = 0.1, *σ* _I_ *→ ∞*. Other parameters are as in Table 1. Each consumer is assumed to ‘occupy’ the space within its exploitation radius, and consumer density is the percentage of total area occupied by all consumers without overlap).

### 3.2 Increasing consumer density leads to social inequality

When consumer density is low, all consumers can adopt an ‘affluent’ harvest-intensively-disperse-far strategy. At low consumer density, everyone can harvest at high rates without causing the resource to deplete globally. However, as consumer density increases, harvesting rate and dispersal radius evolve to lower values (Fig. 2). As consumer density increases further, the intensive harvesting strategy is no longer sustainable, because resources begin to deplete. At this stage, instead of lowering their harvesting rates to adjust to the changed circumstances, some consumers begin to harvest even more intensely to sustain dispersal, while others adopt a ‘frugal’ harvest-prudently-don’t-disperse strategy, by accepting lower resource extraction along with reduced dispersal costs. The proportion of sedentary consumers continues to increase with consumer density, and finally, when space gets overcrowded, all consumers adopt a sedentary strategy.

At all consumer densities, the average evolved harvesting rates (white line in Fig. 2) are higher than the corresponding optimal rates, i.e., the yield-maximizing rate (green) or the profit-maximizing rate (magenta). Consequently, the average resource extraction rate is less than the optimal values. Thus, when left to themselves, consumers over-exploit the resource to some degree and the question arises, when do consumers adapt a yield-maximizing strategy?

### 3.3 Mobile consumers achieve maximum yield at the ‘edge of inequality’, unless dispersal is very cheap

When the benefits of harvesting are low, the population is monomorphic (or ‘sedentary’ regime; region S in Fig. 3A,C), and all individuals have a low harvesting rate (Fig. 3B) and near-zero dispersal (Fig. 3D). The evolved harvesting rates are close to the yield-maximizing rate for sedentary consumers (see Methods). In this regime, resource harvesting is inefficient: the harvesting rate is low (Fig. 4A) despite an abundance of resources in the environment (Fig. 4B). This is because high dispersal costs force consumers to adapt a sedentary lifestyle and prevent them from exploring other resource rich areas. However, since each consumer only harvests resource from his/her own location, there is no competition for resource, and resource distribution is equal.

**Figure 3:**
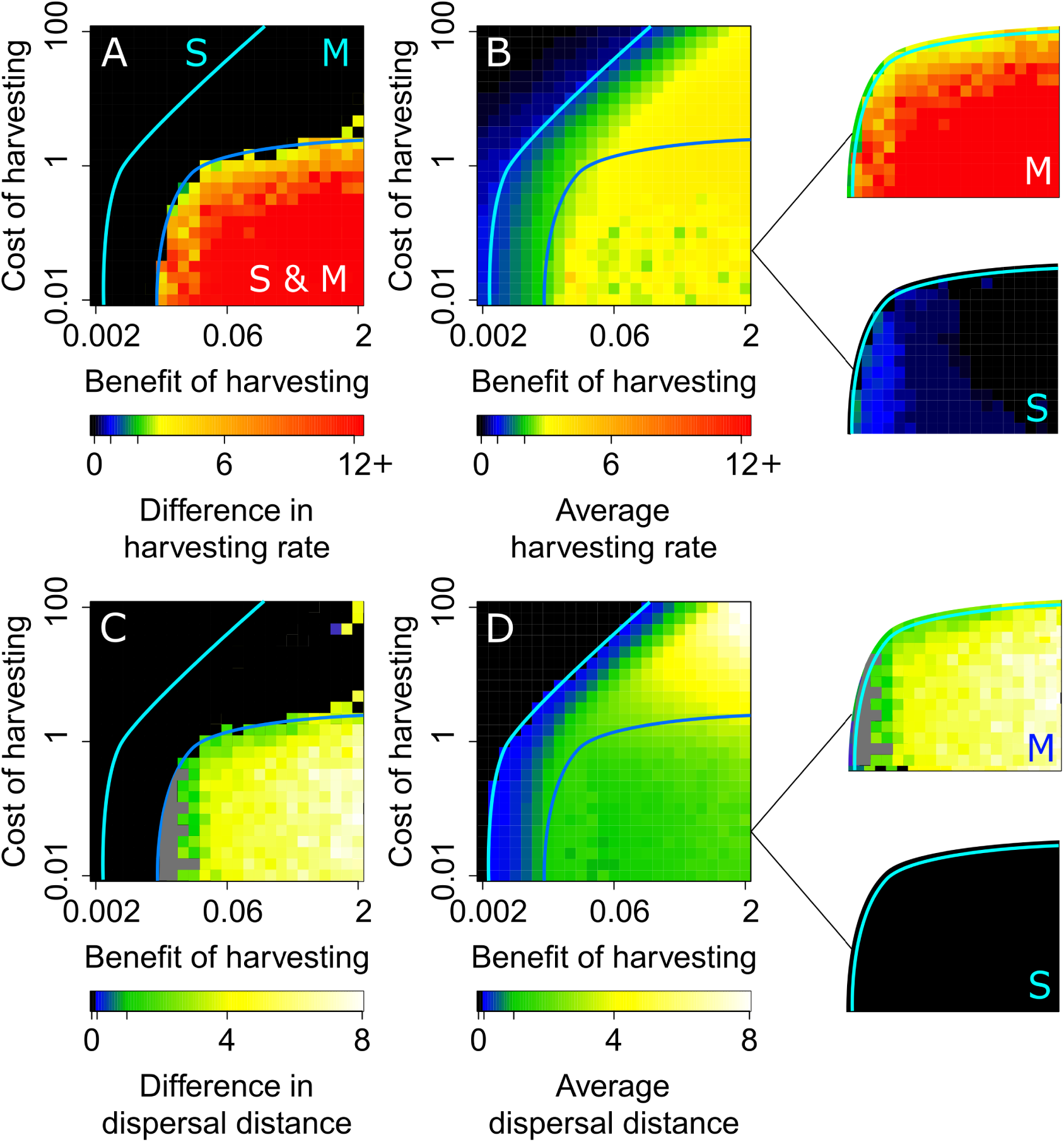
Three regimes of strategies are observed when varying the costs and benefits of harvesting. The difference between the harvesting rate (A) and dispersal distance (C) of sedentary and mobile consumers; and the average harvesting rate (B) and dispersal distance (D) of the population. In A-B, blue and green represent the sedentary and mobile yield-maximizing strategies, respectively (see Methods). Grey areas are where we could not correctly detect a difference (due to insufficient resolution). We find three regimes of strategies (marked by cyan and blue lines): Sedentary (S), in which all consumers are prudent and sedentary, Mobile (M), in which all consumers are overexploitative and mobile, and Coexistence (S & M). For this regime, outsets show the values separately for sedentary and mobile consumers. Parameters: *r* _I_ = 0.1, *σ* _I_ *→ ∞*. Other parameters are as in Table 1.

**Figure 4:**
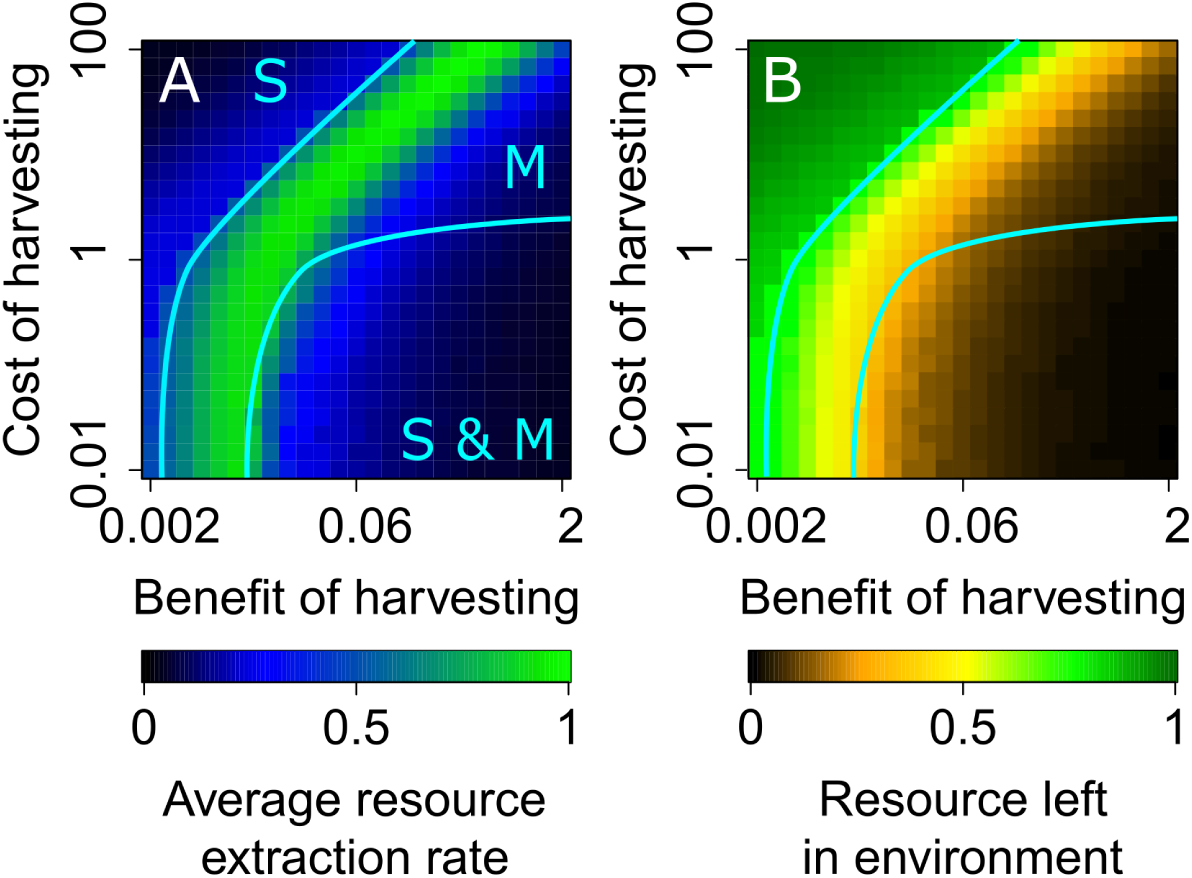
Maximum sustainable resource extraction occurs in the mobile regime. Per capita average resource extraction rate as a fraction of the yield-maximizing rate (A) and the resource left in the environment as a fraction of the carrying capacity (B). The sedentary regime is equitable, but inefficient, because the resource extraction is less than optimal even though there are ample resources in the environment. The mobile regime is both equitable as well as efficient, and the total resource extraction rate reaches its maximum evolved value in this regime. However, for very low dispersal costs, the tragedy of the commons occurs in this regime. The coexistence regime is neither equitable nor efficient. In this regime, the resource extraction rate is suboptimal, and overexploitation by mobile consumers results in the tragedy of the commons.

As the harvesting benefits increase and costs of dispersal decrease, dispersal becomes viable. The population still remains monomorphic, but all consumers now adapt a mobile (harvest and disperse) strategy (‘mobile’ regime; region M in Fig. 3D). The harvesting rate and resource extraction rate both increase, until maximum yield is reached (green band in Fig. 3B and Fig. 4A). At this extraction rate, consumers locally deplete the resource, but the depletion is temporary and resource extraction is globally sustainable. Resource distribution is fair, and all consumers harvest the resource at similar rates. Therefore, resource extraction is efficient as well as fair. However, when both benefits and costs of harvesting are very high (i.e, equivalently, costs of dispersal are very low), the temptation to rapidly harvest the resource is strong, and also feasible as dispersal is cheap. Consequently, consumers begin to overexploit the resource, and even though the population remains monomorphic (everyone evolves high dispersal), resource extraction drops below the optimal value as a consequence of the tragedy of the commons (top right corner in Figs. 3 and 4).

When costs of dispersal are significant, increasing hasvesting benefits lead to a diversification of strategies (‘coexistence’ regime; region S & M). Initially, mobile consumers tend to harvest at high rates. But as the amount of resource in the environment reduces because of overexploitation by these consumers, the sedentary strategy also becomes viable. This is because the benefit forgone by a reduction in harvesting rate is balanced by avoiding the cost of dispersal. As some consumers become sedentary, the density of mobile consumers decreases, allowing them to sustain higher harvesting rates. Therefore, prudent sedentary consumers (cooperators) and overexploitative mobile consumers (cheaters) coexist in the population (Fig. 3A,C). This regime is characterized by a social divide, in which the resource distribution is unequal. Low costs of harvesting allow cheaters to greatly increase their harvesting rates (outsets in 3B). A small number of cheaters thrive at the expense of a large fraction of cooperators (SI-Fig. 5). For lower efficiency values, sedentary consumers adapt the profit-maximizing harvesting rate. However, as efficiency increases further, mobile consumers become increasingly overexploitative, destroying even the neighborhoods of sedentary consumers and driving their resource extraction rates to zero (Fig. 4B, outsets). In this regime, resource distribution may be equal on average over a long time period as each consumer adapts either strategy at different times, but in the short term, this regime is inequitable.

**Figure 5:**
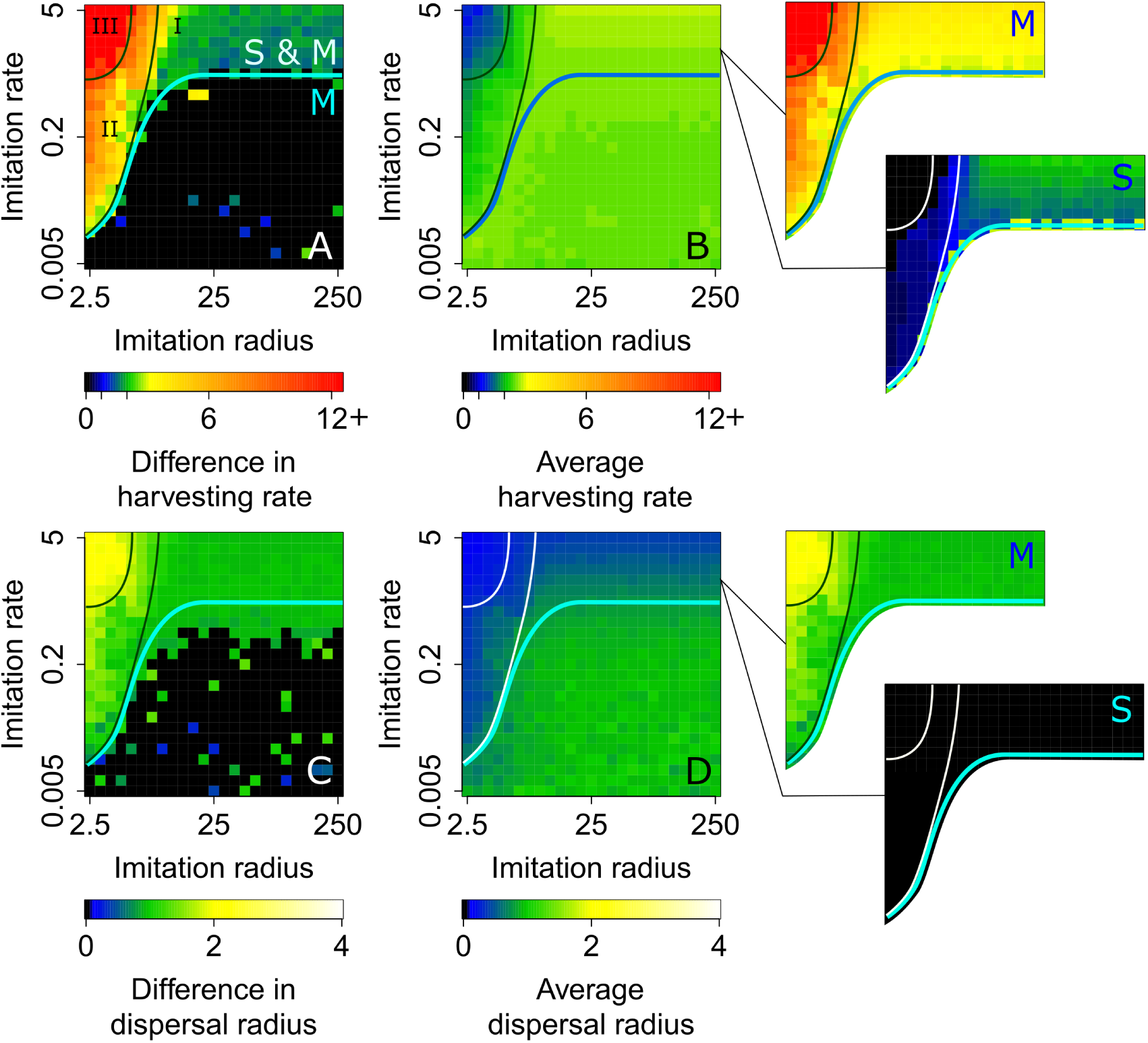
Four regimes of strategies are observed when varying the spatio-temporal scales of imitation. In region M, characterized by low imitation rates, the population is monomorphic with an efficient, mobile strategy. However, as imitation rate increases, the strategy diversifies, taking the system into the coexistence regime (region S & M). We subdivide the S & M region into three regions. In region I, characterized by global information, both sedentary and mobile consumers have high harvesting rates, resulting in the tragedy of the commons. As the imitation radius decreases (region II), (mobile) cheaters become more exploitative, and (sedentary) cooperators harvest at rates close to the sedentary profit-maximizing rate. With even further decrease in imitation radius (region III), cheaters become extremely overexploitative. For more analysis of regions I-III, see SI-Fig. 7. Parameters: *b* = 0.02, *c* _H_ = 0.8. Other parameters are as in Table 1.

### 3.4 Rapid consumer adaptation and localized information both aggravate social inequality

The imitation of strategies by consumers depends on their knowledge of strategies of other consumers. If information on others’ strategies travels far, each consumer has a larger pool of potential strategies to compare with its own and imitate. So far, we assumed that all individuals can sample the entire population for social learning (*σ* _I_ → ∞). We now relax this assumption. Figure 5 shows the evolved strategies as a function of the imitation radius and the imitation rate, for payoff parameters from the mobile regime (See SI-Fig. 6 for a similar analysis in the coexistence regime).

**Figure 6:**
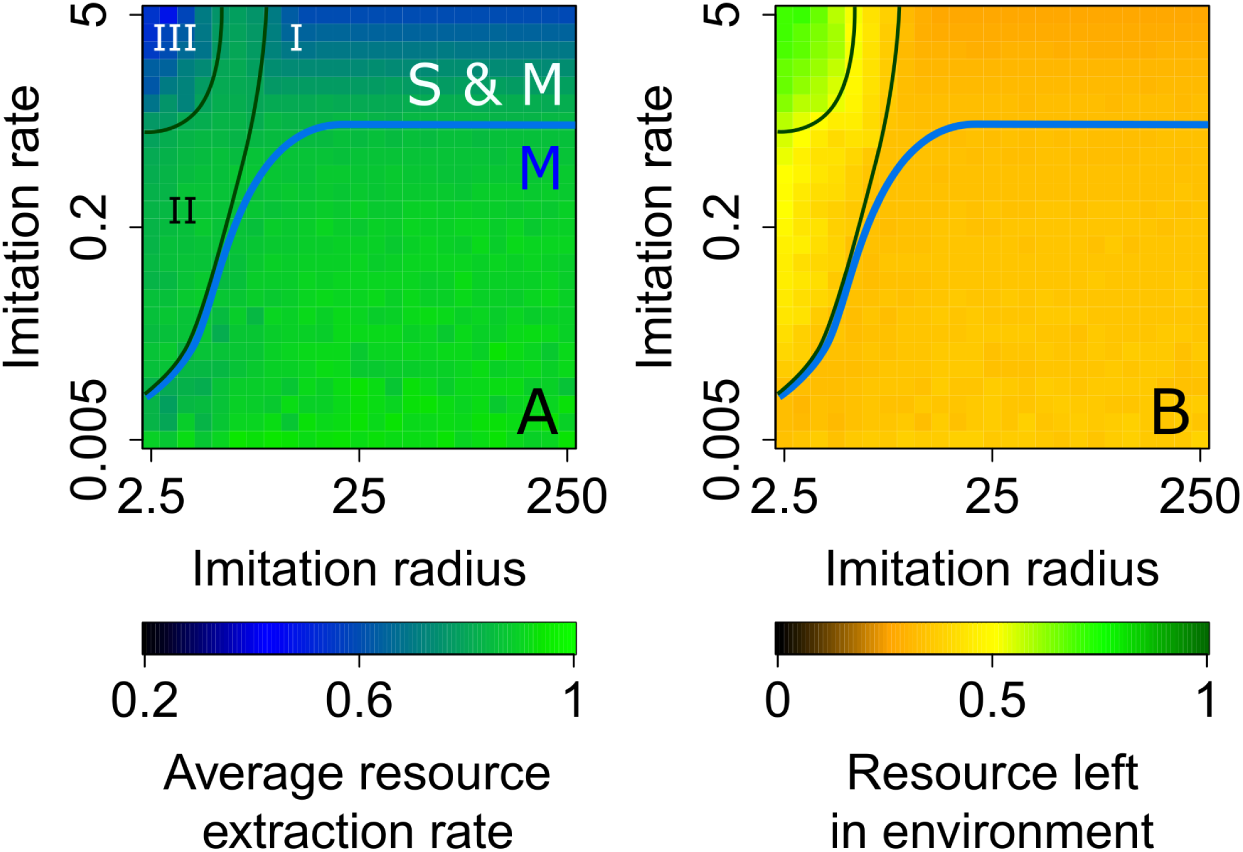
Impatience and myopia among consumers aggravate social inequality. Average resource extraction rate decreases with increasing imitation rate. In region III, this is because only a small fraction of cheaters exploit resource at a high rate, while cooperators, which form a vast majority of the population, get zero resource. In region I, both cooperators and cheaters harvest aggressively and overexploit the resource. Region II allows a substantial number of cheaters to coexist with cooperators. See SI-Fig. 7 for additional analysis of regions I-III.

For low to moderate imitation rates (region marked ‘M’ in Fig. 5A) the population is monomorphic as in the mobile regime, and the resource extraction rate is high (region ‘M’ in Fig. 6). As the imitation rate increases, i.e., as consumers become ‘impatient’, consumers diversify into mobile and sedentary strategies. Fast imitation causes a strategy to change even before its consequences are reflected in the payoffs. Particularly, it prevents sedentary consumers from realizing the long-term benefits of low harvesting rates, leading to an unsustainable increase in harvesting (outset in Fig. 5 and SI-Fig. 7). For large imitation radius (region I), the difference between the harvesting rates of sedentary and mobile consumers is small (SI-Fig. 7). Since all consumers have high harvesting rates, the resource is overexploited (Fig. 6). As a consequence, the cheaters are only marginally better off than cooperators. The cheater strategy becomes more rewarding when consumers are impatient as well as myopic (region III in Fig. 5A). In this regime, cheaters become extremely overexploitative with very high harvesting rates. However, the proportion of cheaters is low (SI-Fig. 7). Therefore this regime is marked by stark social divide, with a handful of cheaters driving extraction rates of the majority of consumers to zero. Resource extraction is acutely inefficient, with much of the resource being left unharvested (Fig. 6). For intermediate spatio-temporal scales of imitation (region II), cheaters are less aggressive, but greater in number. Cooperator harvesting rates are close to the sedentary profit-maximizing value. This leads to greater resource extraction, almost entirely by cheaters (Fig. 6, SI-Fig. 7). In other words, this region is most favourable for cheaters.

### 3.5 Effect of parameters and model variations

In SI-Section S3, we investigate the effect of variations in the remaining model parameters as well as some simple model extensions.

## 4 Discussion

### 4.1 Comparison with previous studies

Spatial public goods games have frequently been used to study the evolution of cooperation. Often, models of spatial public goods rely on additional mechanisms to stabilize cooperation, such as volunteering [34], rewarding cooperators or punishing defectors [28, 50, 51], and conditional strategies [52]. Other models rely on a spatial structure (inherent or emergent) that support clustering of cooperators [35, 53, 54]. In all these studies cooperation has been defined in terms of game-theoretic payoffs, and whether an act is coperative or defective is independent of the socio-environmental conditions. By contrast, cooperation and defection in our model can only be defined in relation to the resource: cooperation is the strategy that ensures that total resource extraction rate is less than the maximum sustainable yield.

When movement is costly, consumers of a spatial resource may face a ‘milker-killer dilemma’ [55, 56], in which each consumer has a choice to be a milker (like our sedentary consumers) or a killer (like our mobile consumers). Several studies have tried to identify the conditions under which either milker and killer strategies are favored [55, 57, 58]. But none of these studies had found coexistence of milkers and killers. Recently, [45] showed a coexistence of sessile cooperators (like milkers) and mobile defectors (like killers) by allowing the coevolution of cooperation and dispersal. In agreement with their results, we found emergence and coexistence of prudent sedentary consumers and mobile overexploitative consumers. However, in our model, direct interactions between individuals are absent, and payoffs of cooperation and dispersal are realized only through the resource. When cheaters begin to overexploit the resource, the incentive to overexploit diminishes and the sedentary, prudent harvesting strategy becomes equally attractive. Therefore, resource dynamics converts cooperation into a beneficial strategy and allows cooperation to be maintained.

Another recent study by [44] used an infinite island model to study the coevolution of cooperation (in the form of investments to public goods) and dispersal (in the form of the rate of leaving the current patch). They found that both dispersal and cooperation are favoured if catastrophes wipe out local populations at an intermediate rate. This is because catastrophes reduce local population densities and increase relatedness among the remaining individuals. It would be interesting to examine the effect of catastrophes in our model. On the one hand, catastrophes may prevent strategy diversification, because a decrease in consumer density brought about by a catastrophe may allow all consumers to sustain high harvesting rates. On the other hand, catastrophes may facilitate coexistence due to the well-known competition-colonization tradeoff.

### 4.2 Extensions and further directions

Our model makes deliberately simple assumptions to capture the essential processes governing strategy evolution. We have assumed that all consumers have the capacity to adapt any strategy they wish. Thus, a sedentary consumer with very low resource extraction rate can switch to an expensive mobile strategy. Because of frequent switching of strategies, all individuals in our population have equal payoffs in the long term, even when two distinct strategies exist at the population level. In real situations however, resource-poor individuals may not be able to afford a strategy switch, forcing them permanently into a low extraction strategy. These effects can be further investigated by incorporating a cost of switching strategies.

Although we have investigated the dependence of evolved strategies on population density, we have kept the population size constant. However, it is well known that increased resource consumption resulting from higher efficiency and better technology leads to a rapid increase in population size. This in turn may necessitate further increase in efficiency. It is possible to incorporate such feedbacks in our model. Exploring the fate of a society in the presence of such feedbacks would be an interesting direction for further research.

## 5 Conclusion

In conclusion, we have explored how dynamics of a renewable resource influence the harvesting and dispersal strategies of consumers. We found that increased consumer density, increased efficiency of resource harvest, reduced costs of dispersal, rapid adaptation by consumers, and localized information on others, are features that lead to a spontaneous diversification of consumers into prudent sedentary consumers and overexploitative mobile ones. This social divide is always accompanied by overexploitation of resources. In modern societies, developments such as technological progress have led to an increase in resource harvesting efficiency and reduced costs of dispersal. This has in turn led to an increase in population size. Furthermore, the pace of life has increased, which in our model, reflects in the pace of strategy adaptation. While these are positive developments as such, our model suggests that they may pose a threat to social equality and sustainability of resource. Our model also suggests a way of mitigating these threats: increasing information availability, which is also a characteristic feature of modern societies. We hope that further work can build on the findings presented here and shed light on the foundations of human and animal behaviour and its changes in response to changing socio-ecological conditions.

## 6 Acknowledgements

The authors would like to thank Vishwesha Guttal for providing computational facilities (via DST-FIST and DBT-IISc partnership programme). JJ would like to thank TIFAC, Government of India, for providing travel support and stipend for carrying out this work at IIASA, Austria. He would also like to thank MHRD, Government of India, for his PhD scholarship. ÅB gratefully acknowledges support from the Swedish Research Council.

## 8 Author contributions

JJ, ÅB, and UD designed the study; JJ performed the simulations; JJ, ÅB, and UD analysed the results and wrote the paper.

